# Mechanisms underneath *Xanthomonas citri* subsp. *citri* promoting of *Agrobacterium-*mediated transient expression in citrus

**DOI:** 10.1101/2023.09.12.557436

**Authors:** Tirtha Lamichhane, Nian Wang

## Abstract

*Agrobacterium-*mediated transient expression (TE) is an important tool in plant genetics study and biotechnology. Citrus is one of the top tree crops, but *Agrobacterium-*mediated TE remains problematic in citrus. Previous study has shown that pretreatment of citrus leaves with *Xanthomonas citri* subsp. *citri* (*Xcc*), which causes citrus canker in almost all citrus cultivars, significantly improves the TE efficacy. Here we have shown that *Xcc* promotes *Agrobacterium-* mediated TE via improving *Agrobacterium* growth, suppressing plant immune response, triggering cell division, and upregulating plant cell wall degrading enzymes. We demonstrate that *Xcc* promotes *Agrobacterium-*mediated TE via PthA and mutation of *pthA4* abolishes the promoting effect of *Xcc.* PthA4 is known to trigger cell division of citrus. We have demonstrated the causative relationship between cell division and *Agrobacterium-*mediated TE using cell division inhibitor mimosine. Importantly, the differential effects of mimosine and colchicine on *Xcc* promoting *Agrobacterium-*mediated TE indicate that S-phase is critical for agroinfiltration of citrus tissues. In addition, PthA4 is known to upregulate plant growth hormones auxin (IAA), gibberellin and cytokinin, as well as cell degrading enzymes (e.g., cellulase). Exogenous application of IAA, cytokinin, and cellulase but not gibberellin significantly improved *Agrobacterium*-mediated TE in leaves of sweet orange, pummelo, Meiwa kumquat, lucky bamboo and rose mallow. Our study provides a mechanistic understanding regarding how *Xcc* promotes *Agrobacterium-*mediated TE and identified two practical measures to improve *Agrobacterium-*mediated TE via pretreatment with plant hormones (i.e., auxin and cytokinin) and cellulase.

## Introduction

*Agrobacterium*-mediated transformation is a widely used tool in plant genetics and biotechnology, which includes both transient and stable expression of genetic information carried by the T-DNA (Hooykaas, 2023). Stable transformation enables the generation of GMO plants with heritable traits (Hooykaas, 2023). However it is a time consuming and labor intensive process (Gelvin, 2005). Transient expression (TE) system is an alternative to the transgenic approach which includes biolistic bombardment, protoplast transient transformation and *Agrobacterium*-mediated transient transformation (Li *et al*., 2009). Leaf agroinfiltration based transient gene expression has become a powerful tool for rapid gene function analysis in plants by virtue of its simplicity and efficiency (Wu *et al*., 2014). It is widely used for rapid assays of gene functions, protein-protein interactions, subcellular localization and CRISPR Cas genome editing (Belhaj *et al*., 2015; Li *et al*., 2009). *Agrobacterium-*mediated TE in plants also offers cost effective and scalable production of plant derived biopharmaceutical products used in human medicine (Chen *et al*., 2014).

*Agrobacterium* is a plant pathogenic bacteria that can infect wide range of eukaryotic cells from fungi to human cells under laboratory conditions (Citovsky *et al*., 2007). Plant species are either susceptible or recalcitrant to *Agrobacterium* infection and the transformation efficiency correlates with the competence between *Agrobacterium* and plant species (Subramoni *et al*., 2014). Higher transformation efficiency is essential to obtain optimal transient expression in plants (Zhang *et al*., 2000). The underlying mechanism of how host plant cells become more competent to *Agrobacterium* transformation is poorly understood (Gelvin, 2021). Many host plant related factors affect the transformation including plant defense, cell division, growth and cell cycle progression stage in transforming host plant cells (Gordon-Kamm *et al*., 2002; Hwang *et al*., 2017; Villemont, *et al*., 1997). For instance, suppression of plant immunity through inducible expression of the bacterial effector AvrPto or transgenic expression of *NahG* increases *Agrobacterium-*mediated TE (Rosas-Díaz *et al*., 2017; Tsuda *et al*., 2012). Host cell proliferation and cell cycle dynamics are critical in plant cell transformation. It was reported that host cell cycle stage S-phase (DNA synthesis) is absolutely required for *Agrobacterium*-mediated T-DNA transfer in plants (Villemont *et al*.,1997). Similarly another study shows that increased stimulation of cell division and growth in maize has increased transformation efficiency (Gordon-Kamm *et al*., 2002).

Citrus is an economically important fruit crop worldwide. Citrus production faces many biotic and abiotic challenges including drought, heat, and diseases (Jia *et al*., 2022). Genetic study and genetic improvement of citrus are paramount to overcome those challenges. Citrus varieties are incompetent with *Agrobacterium*; therefore, *Agrobacterium-*mediated TE remains problematic in citrus. Interestingly, our previous study has shown that pre-treatment of leaves of multiple citrus genotypes with *Xanthomonas citri* subsp. *citri* (*Xcc*) improves *Agrobacterium*-mediated TE efficiency (Jia and Wang, 2014a). However, the underneath mechanism for the improved *Agrobacterium-*mediated TE enabled by *Xcc* is unknown, which is explored in this study.

## Results

### *Xcc* Promotes *Agrobacterium* Growth in Citrus

*Agrobacterium*-mediated TE assays in plant tissues are reported to peak around 2 - 4 days after inoculation, which predominantly results from the T-DNA not integrated into the host genome (Krenek *et al*., 2015). Therefore, 1 and 4 <u>d</u>ays <u>p</u>ost agroinfiltration (dpi) were selected to investigate the interactions between *Agrobacterium*-*Xcc*-Citrus. We first investigated whether *Xcc* improves *Agrobacterium*-mediated TE by simply promoting the growth of *Agrobacterium* in leaves. *Xcc* had no effect on *Agrobacterium* titers at 1 dpi, but did increase *Agrobacterium* titers at 4 dpi (Fig. 1A). However, *Xcc* only increased *Agrobacterium* titers approximately 3.5% and 4.0% compared to the water treatment and no treatment, respectively (Fig. 1A). In addition, *Xcc* had no effect on *Agrobacterium* growth on NA plates (Supplementary Fig. 1) and in the non-host plant *Nicotiana benthamiana* (Fig. 1B), indicating that the physiological and cellular changes of citrus leaves resulting from *Xcc* infection are responsible for the growth promotion of *Agrobacterium* at 4 dpi.

**Figure 1.**
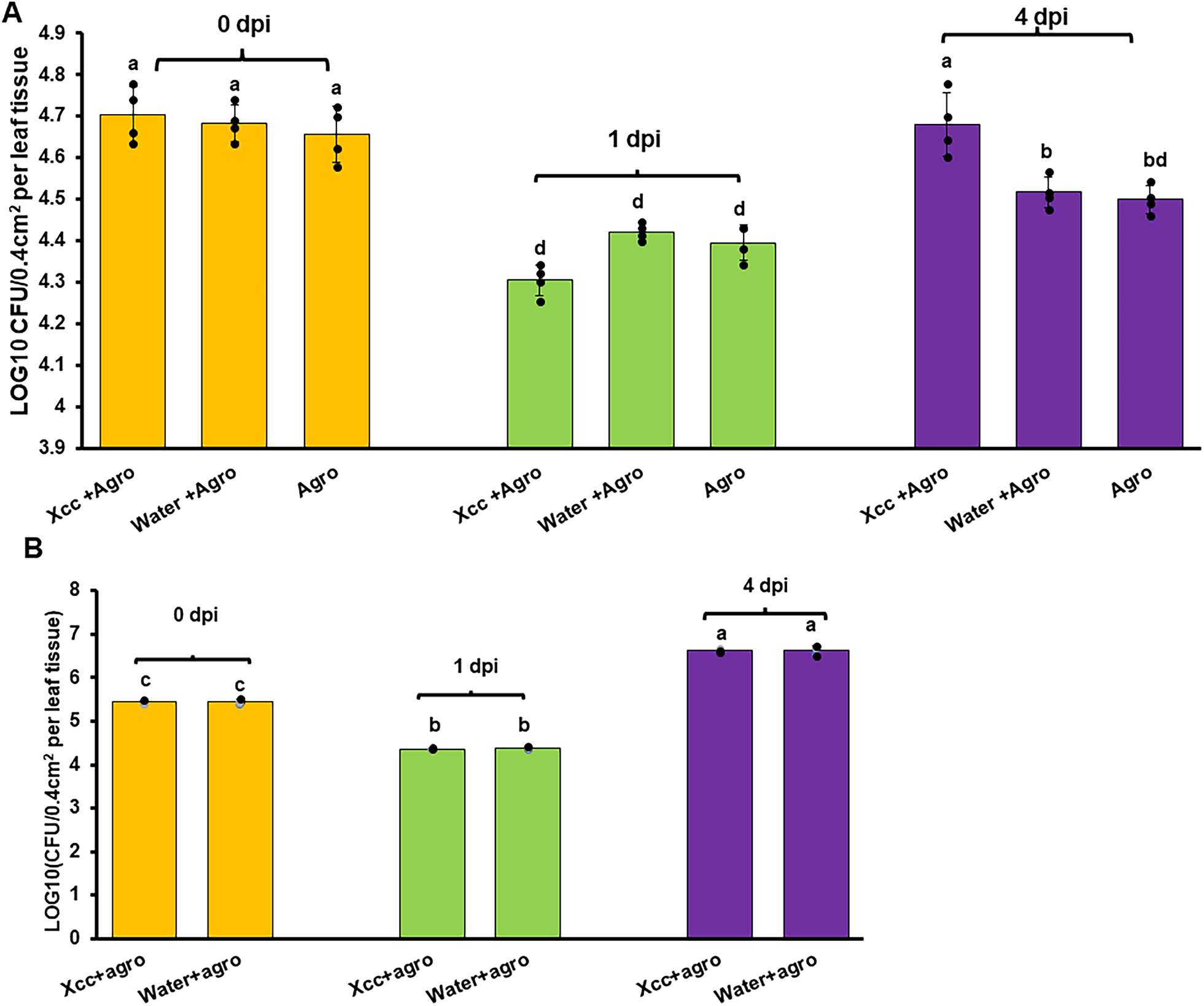
*Agrobacterium tumefaciens* growth in agroinfiltration assay in sweet orange (*C. sinensis*) cv. Hamlin (A) and in *Nicotiana benthamiana* (B). *Agrobacterium tumefaciens* strain EHA 105 carrying GUS binary vector (OD600 =0.8) was used for agroinfiltration. 0 dpi indicates the time point of sample collection within one hour of agroinfiltration. Citrus and tobacco leaves were treated with *Xanthomonas citri* subsp. *citri* (Xcc, 5 X 10^8^ CFU/ml) or water 12 hours before agroinfiltration. The values represent mean ± SD (n=4). Different letters indicate significant difference (P<0.05). For A, Statistical analysis was conducted by repeated measures analysis of variance (ANOVA) using the MIXED procedure in SAS V9.4 (SAS Institute Inc). Evaluated main effects include bacterial treatment and time. For B, one-way ANOVA with post-hoc Tukey Test was conducted. Agro: Agroinfiltration; DPI: days post agroinfiltration. The experiments were conducted at least twice with similar results.

### *Xcc* Effect on Plant Immune Response

Next, we investigated whether *Xcc* suppresses plant immune responses during Agroinfiltration. At 4 dpi, accordant with the increased *Agrobacterium* titers, *Xcc* decreased expression of *PR1*, *PR2* and *PR5*, which are marker genes of plant immune responses, compared to water treatment (Fig. 2). Previous research indicated that constitutive expression of *PR* genes (*PR1*, *PR2* and *PR5*) in *Arabidopsis* reduces susceptibility to *Agrobacterium* infection (Veena *et al*., 2003). On the other hand, at 1 dpi, *Xcc* preinfection was associated with higher expression of *PR1*, but had no effect on *PR2* and *PR5* compared to water treatment (Fig. 2) which was consistent with the similar *Agrobacterium* titers at 1 dpi for the *Xcc* and water treatments. Reactive oxygen species (ROS) play important roles in plant immune responses against pathogens and can also have direct antimicrobial effect against pathogens (Jones and Dangl, 2006). ROS levels were similar for all the treatments at both 1 and 4 dpi (Supplementary Fig. 2), indicating that ROS do not play a significant role in *Xcc* promoted *Agrobacterium* growth and *Agrobacterium*-mediated TE at 4 dpi.

**Figure 2.**
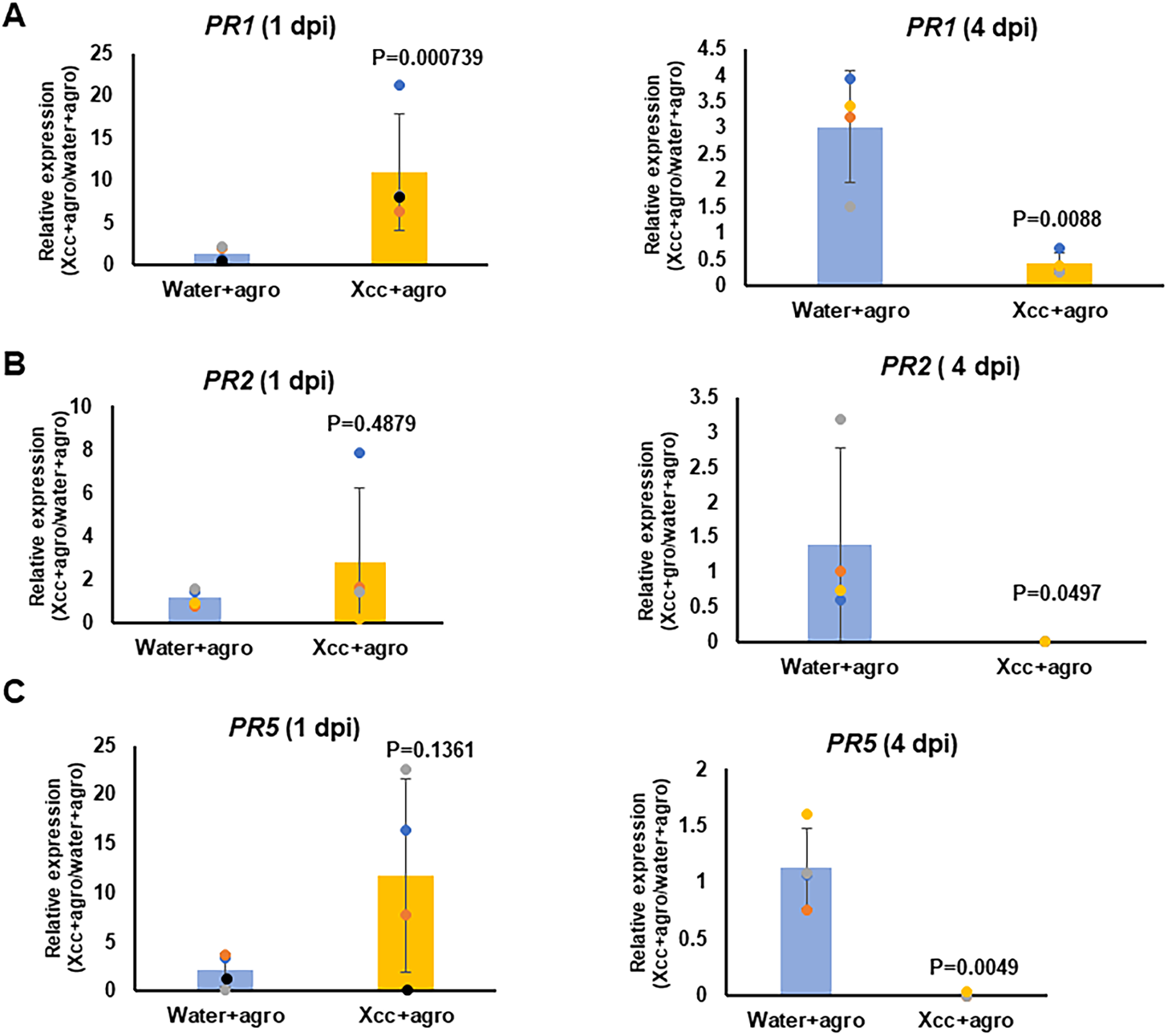
Expression of pathogenesis-related (*PR*) genes in agroinfiltration of citrus via RT-qPCR. Citrus leaves were treated with *Xcc* of 5X10^8^ CFU/ml and water, followed by agroinfiltration after 12 hours. *Agrobacterium tumefaciens* strain EHA 105 (OD_600_=0.8) carrying the GUS binary vector was used for agroinfiltration. DPI: days post inoculation by *Agrobacterium*. 0 dpi: the time point of sample collection within one hour of agroinfiltration. Citrus *GAPDH* gene was used as an internal control for normalization. A. *PR1*; B. *PR2*. C. *PR5*. Error bars indicate mean ± SD of four independent biological replicates. Student *t* test was conducted for statistical analysis. The experiment was repeated at least twice with comparable results.

### PthA4 Triggered Cell Division Plays Critical Roles in *Xcc* Promoting *Agrobacterium-* Mediated TE

Plant cells undergoing active cell divisions are prone to *Agrobacterium-*mediated transformation (Gordon-Kamm *et al*., 2002). *Xcc* is known to cause excessive cell division and enlargement of citrus cells of mesophyll tissues via PthA4 (Duan *et al*., 2018; Duan *et al*., 1999; Pereira *et al*., 2014). Our previous study showed that PthA4 is required for the *Xcc*-facilitated *Agrobacterium-*mediated TE in grapefruit (*Citrus paradisi*) (Jia and Wang, 2014a). Mutation of *pthA4* abolished *Xcc* promotion of *Agrobacterium-*mediated transient expression in *C. sinensis* based on both GUS and GFP assays (Fig. 3, Supplementary Figs. 3 and 4). CDKA kinase belongs to cyclins and cyclin-dependent protein kinase family (CDK) and plays key roles in cell cycle progression and is required for G1/S transition (Gutierrez, 2009), whereas CDKBs, another class of CDK, function in mitotic entry from G2 to M transition (Boudolf *et al*., 2004). The expression of cell division related genes (*CDKA* (*Cs6g19580)*, *CDKB1-2* (*Cs6g10590*) and *CDKB2-2* (*Cs3g27820*)) was reduced in the leaves treated with the *pthA4* mutant compared to the wild type *Xcc* and the complementation strain (Fig. 4).

**Figure 3.**
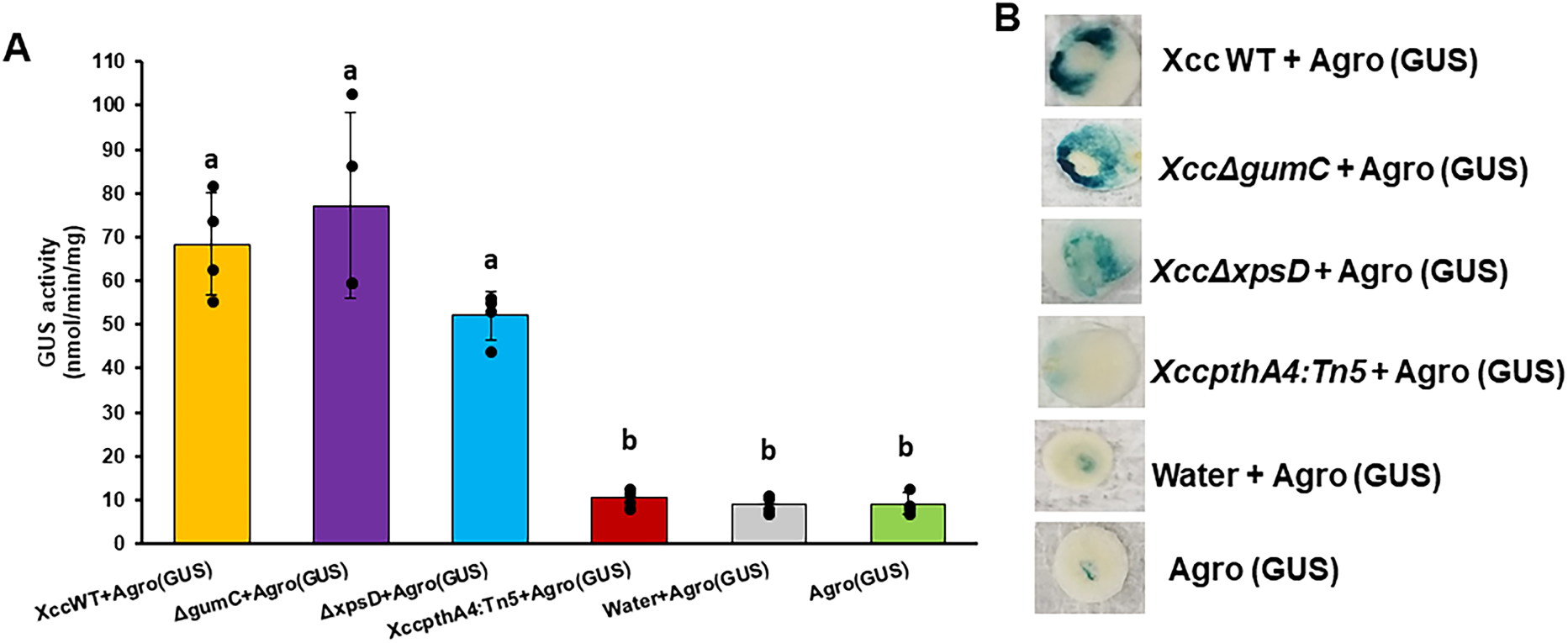
Citrus leaves agroinfiltration assay with different Xcc strains. A. GUS assay. Citrus leaves were treated with 5X10^8^ CFU/ml of Xcc strains 12 hours before agroinfiltration. *Agrobacterium tumefaciens* strain EHA 105 carrying GUS binary vector was used for agroinfiltration. GUS assay was performed at 4 dpi. GUS activity was measured by collecting leaf discs from agroinfiltrated leaf area. The values represent mean ±SD (n =4). Different letters indicate significant difference (P<0.05, One-way ANOVA with post-hoc Tukey Test). Before analysis, the data were checked for normality and homoscedasticity using Shapiro-Wilk test and Levene’s test. respectively, and were log10 transformed to satisfy assumptions of normality and homoscedasticity. Mean separation was made based on Tukey test (a = 0.05) with the log10 transformed data. All experiments were repeated twice with similar results. Agro: Agroinfiltration; DPI: days post agroinfiltration; *Xcc*: *Xanthomonas citri* subsp. *citri*; WT: wild type; ΔgumC: *gumC* mutant of *Xcc*; ΔxpsD: *xpsD* mutant of *Xcc*; XccpthA4:Tn5: *pthA4* Tn5 insertion mutant of *Xcc*. Water treatment and agroinfiltration alone were used as negative controls. B. Histochemical GUS staining of citrus leaf tissues in A. Leaf discs were stained overnight in X-gluc substrate at 37 °C and chlorophyll was removed by ethanol (70%) and photographs were taken.

**Figure 4.**
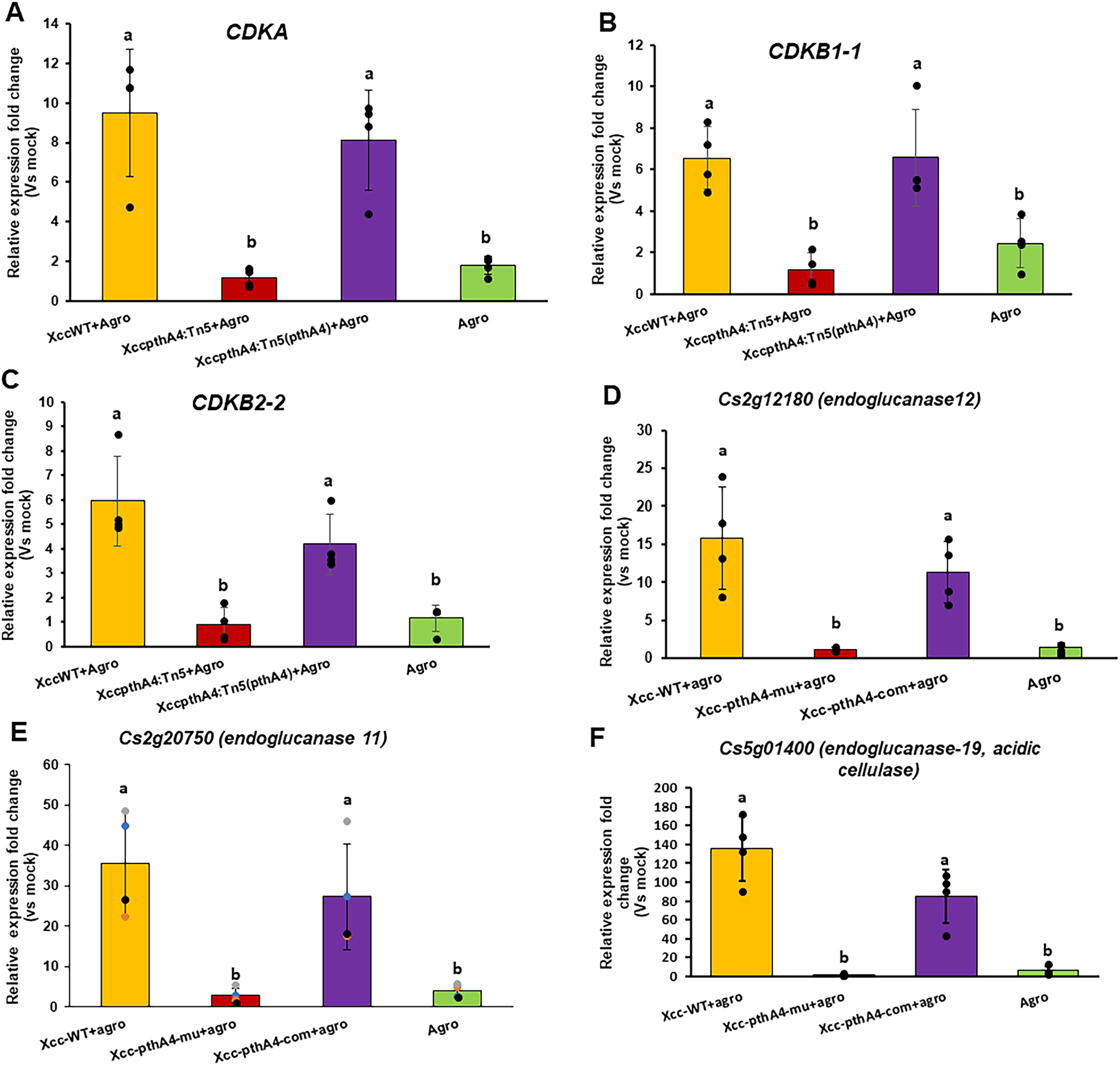
Expression of genes related to cell division and cell wall degrading enzymes in agroinfiltration assay of citrus. Citrus leaves were treated with 5X10^8^ CFU/ml of *Xcc* strains (WT, *pthA4* mutant and *pthA4* mutant complementation strain) followed by agroinfiltration 12 hours after treatment. Agroinfiltration was performed with *Agrobacterium tumefaciens* strain EHA 105 carrying the GUS binary vector. Agroinfiltration without any treatment was performed as a negative control (Agro). RT-qPCR was conducted 4 days post agroinfiltration. Pretreatment with water was used as the mock inoculation control. Citrus *GAPDH* gene was used as internal control for normalization. **A, B** and **C** show expression of cell division marker genes *CDKA*, *CDKB1-1* and *CDKB2-2,* respectively. **D, E, F** represent expression of citrus endoglucanase genes. Error bars indicate mean ±SD (n=4). One way ANOVA followed by Tukey HSD test was conducted for statistical analysis. Different letters indicate significant statistical difference (p<0.05).

To further investigate how cell division affects *Xcc*-facilitated *Agrobacterium-*mediated TE, we tested two different cell division inhibitors: mimosine and colchicine. Both 1 mM and 10 mM mimosine significantly reduced the *Agrobacterium-*mediated TE in *C. sinensis* compared to the control based on GUS assays (Fig. 5), but colchicine at 1 mM and 10 mM had no inhibitory effect (Fig. 5). Mimosine blocks cell cycle progression before S-phase entry while colchicine causes metaphase arrest during the M phase (Planchais *et al*., 2000). This is in accordance with the previous finding that S-Phase is critical in T-DNA transfer (Villemont *et al*., 1997).

**Figure 5.**
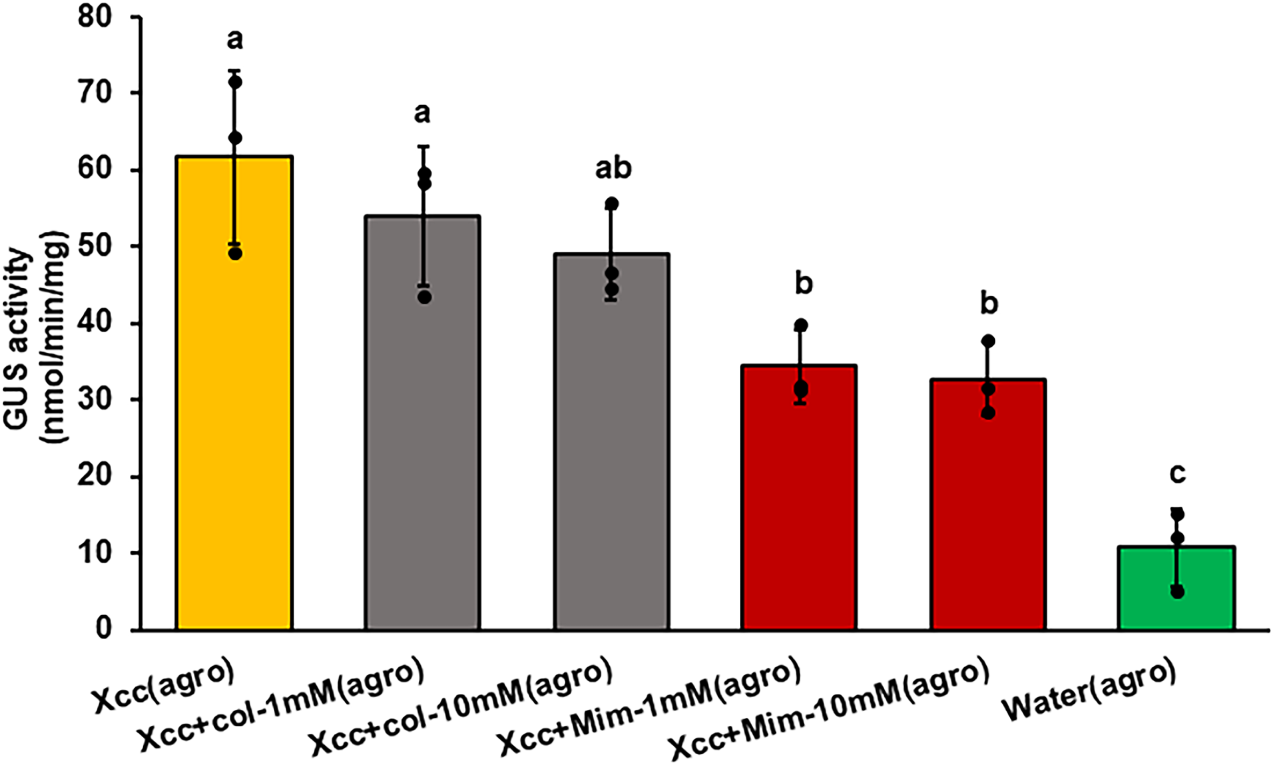
Effect of cell cycle inhibitors on *Xcc-facilitated* agroinfiltration efficiency in citrus. *Agrobacterium tumefaciens* strain EHA105 (OD_600_=0.8) carrying the GUS binary vector was used in agroinfiltration assay. Citrus leaves were treated with *Xcc* WT (5X10^8^ CFU/ml) and cell cycle inhibitors (1:1 (v:v)) and followed by agroinfiltration on the same treatment site after 12 hours. Water treatment followed by agroinfiltration was used as a negative control. GUS activity was measured 4 days post agroinfiltration. Error bars represent mean ± SD (n =4). All experiments were repeated twice with similar results. One way ANOVA with Tukey HSD test was conducted for statistical analysis. Different letters indicate significant statistical difference (p<0.05). col: colchicine. Mim: mimosine.

### Auxin and Cytokinin but not Gibberellin Promote *Agrobacterium-*Mediated TE

PthA4 is known to up-regulate auxin and gibberellin response genes (Pereira *et al*., 2014). PthA4 also positively regulates cytokinin metabolism via its target gene *LOB1* in citrus (Hu *et al*., 2014; Zou *et al*., 2021). Auxin, gibberellin and cytokinin positively regulate cell division (Shimotohno *et al*., 2021). We tested whether plant hormones that positively regulate cell division promote *Agrobacterium-*mediated TE. Indole-3-acetic acid (IAA) of 10 mM and 20 mM, but not 1 mM significantly increased *Agrobacterium-*mediated TE in *C. sinensis* compared to the negative control based on GUS assays (Fig. 6). Kinetin (a cytokinin-like synthetic plant hormone) of 25 mM and 50 mM, but not 5 mM also significantly boosted *Agrobacterium-*mediated TE (Fig. 6). Gibberellin (5 mM and 50 mM) had no effect on *Agrobacterium-*mediated TE (Fig. 6).

**Figure 6.**
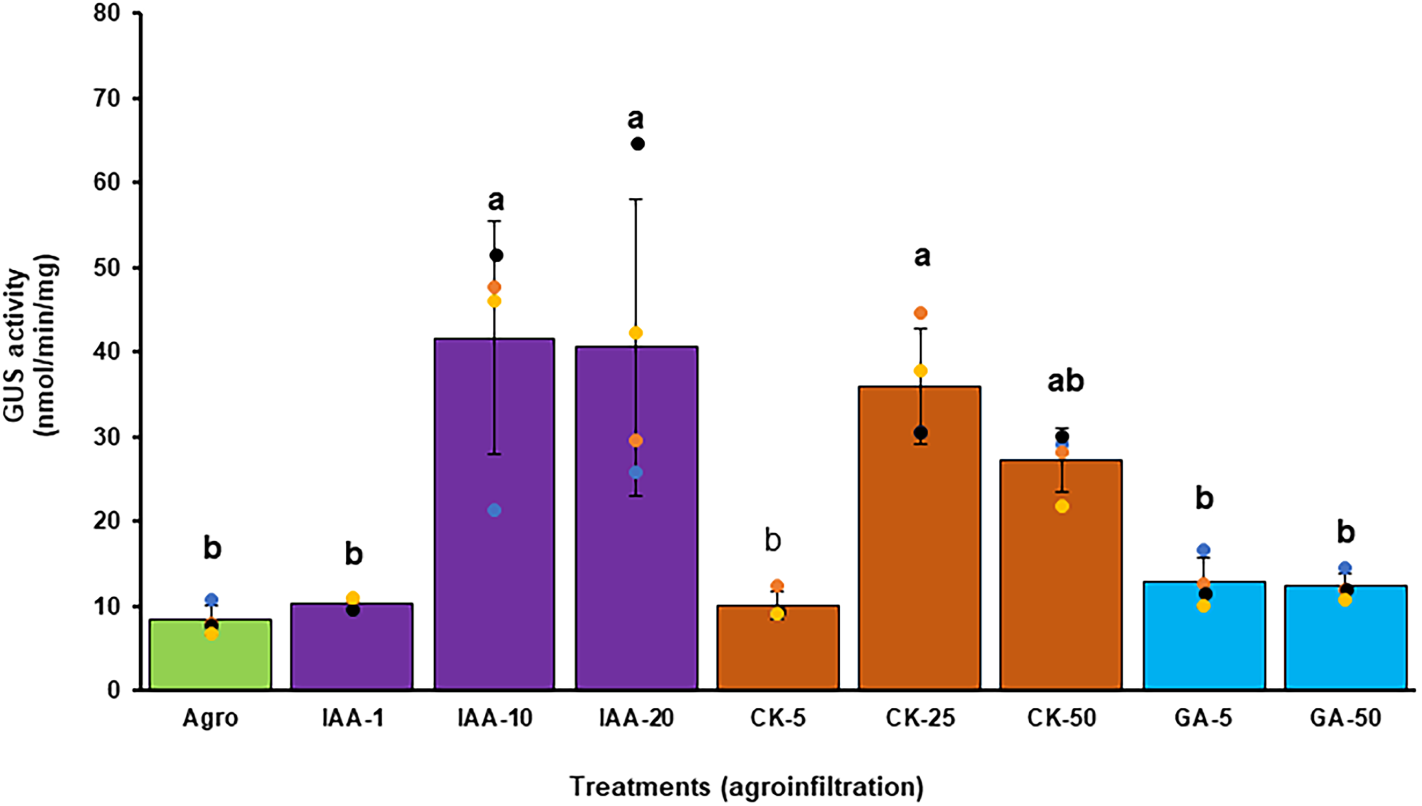
Effect of plant growth phytohormones on agroinfiltration efficiency in citrus. *Agrobacterium tumefaciens* strain EHA105 (OD_600_=0.8) carrying the GUS binary vector was used for agroinfiltration. Citrus leaves were first treated with IAA (1 mM, 10 mM, and 20 mM), kinetin (a cytokinin-like synthetic plant hormone, 5 mM, 25 mM, and 50 mM), and gibberellin (5 mM and 50 mM) and followed by agroinfiltration on the same treatment site after 12 hours. Agroinfiltration alone without any treatment was used as a negative control (Agro). GUS activity was measured 4 days post agroinfiltration. Error bars represent mean ± SD (n =4). All experiments were repeated twice with similar results. One way ANOVA with Tukey HSD test was conducted for statistical analysis. Different letters indicate significant statistical difference (p<0.05).

### Upregulation of Plant Cell Wall Degrading Enzymes by *Xcc* Promotes *Agrobacterium-* Mediated TE

Because cell division inhibitor mimosine did not completely block *Xcc* promotion of *Agrobacterium*-mediated transient expression, we hypothesized that other factors are involved. Phenolic compounds (such as acetosyringone and hydroxy-acetosyringone) and sugars (such as galactose) are known to induce the expression of *vir* genes and subsequently activate T-DNA transfer (Scott *et al*., 1985; Cangelosi *et al*. 1990). Both phenolic compounds and sugars are components of the plant cell wall (Gangl and Tenhaken, 2016; Gupta and De, 2017). *Xcc* secretes cell wall degrading enzymes like cellulases, xylanases, lipases, and proteases via the type II secretion system (T2SS) (Jalan *et al*., 2011). Surprisingly, knock-out of *xpsD*, which encodes the outer membrane secretin of the T2SS and is indispensable for the secretion of cell wall degrading enzymes (Szczesny *et al*., 2010), had no effect on *Agrobacterium*-mediated TE (Fig. 3, Supplementary Fig. 3). For comparison purposes, we also included the *gumC* mutant (Fig. 3, Supplementary Fig. 3). GumC is required for xanthan gum production and biofilm formation (Li and Wang, 2011; Rigano *et al*., 2007). Disruption of *gumC* had no significant difference from the wild type *Xcc* on *Agrobacterium*-mediated TE (Fig. 3, Supplementary Fig. 3).

We then investigated the involvement of plant cell wall degrading enzymes. PthA4 positively regulated multiple genes encoding endoglucanases including Cs2g12180, Cs2g20750, Cs5g01400, and Cs2g17090 (Fig. 4, Supplementary Fig. 5), suggesting that *Xcc* upregulates plant cell wall degrading enzymes to facilitate *Agrobacterium*-mediated transient expression. Consistent with this notion, cellulase, which includes endoglucanase (Jecu, 2000), significantly increased *Agrobacterium*-mediated TE compared to the negative control based on GUS assay (Fig. 7).

**Fig. 7.**
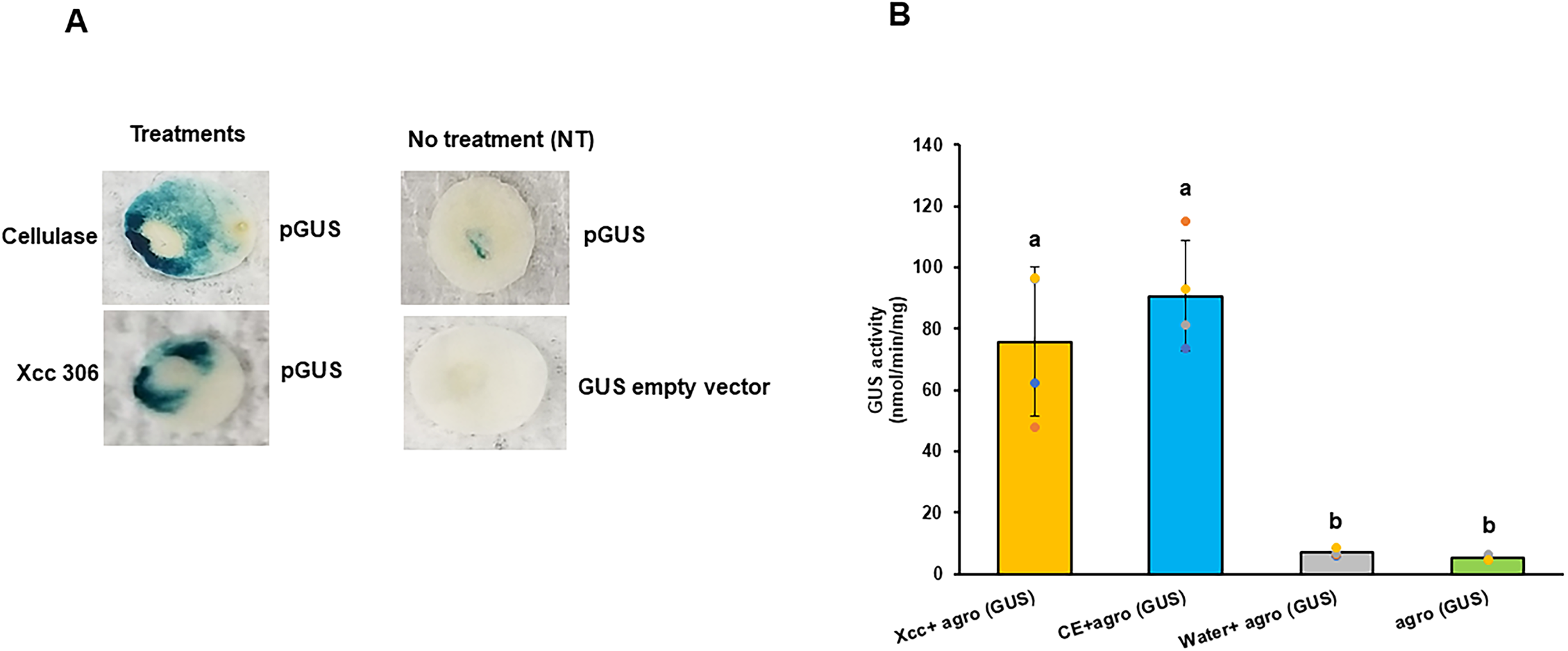
Cellulase treatment enhances *Agrobacterium* mediated transient expression assay in citrus leaves. *Agrobacterium tumefaciens* strain EHA 105 carrying GUS binary vector (OD600 =0.8) was used for agroinfiltration (agro). Citrus leaves were pre-treated with cellulase (CE, 2 g/L) or *Xanthomonas citri* subsp. *citri* (Xcc, 5 X 10^8^ CFU/ml) 12 hours before agroinfiltration. Water treatment or agroinfiltration alone were included as negative controls. A. Histochemical GUS staining of citrus leaves four days post agroinfiltration (dpi). B. GUS assays at 4 dpi. Xcc/enzyme treated GUS empty vector agroinfiltrated citrus leaves were used as controls. Error bars represent mean ± SD (n =4). All experiments were repeated twice with similar results. Different letters indicate significant statistical difference (One way ANOVA followed by Tukey HSD test, p<0.05).

### Cellulase, IAA, and Cytokinin Treatments Enhance *Agrobacterium-*Mediated TE in multiple Plants

Next, we tested whether cellulase, IAA, and cytokinin treatments can improve *Agrobacterium-*mediated TE in other plants. We did not include *Xcc* because of its limited host range on citrus. Cellulase (20 mg/ml), IAA (20 mM), and kinetin (20 mM) treatments all significantly increased GUS activities in leaves of pummelo (*C. maxima*), Meiwa kumquat (*Fortunella crassifolia*), lucky bamboo (*Dracaena sanderiana*) and rose mallow (*Hibiscus rosa-sinensis*) (Fig. 8).

**Fig. 8.**
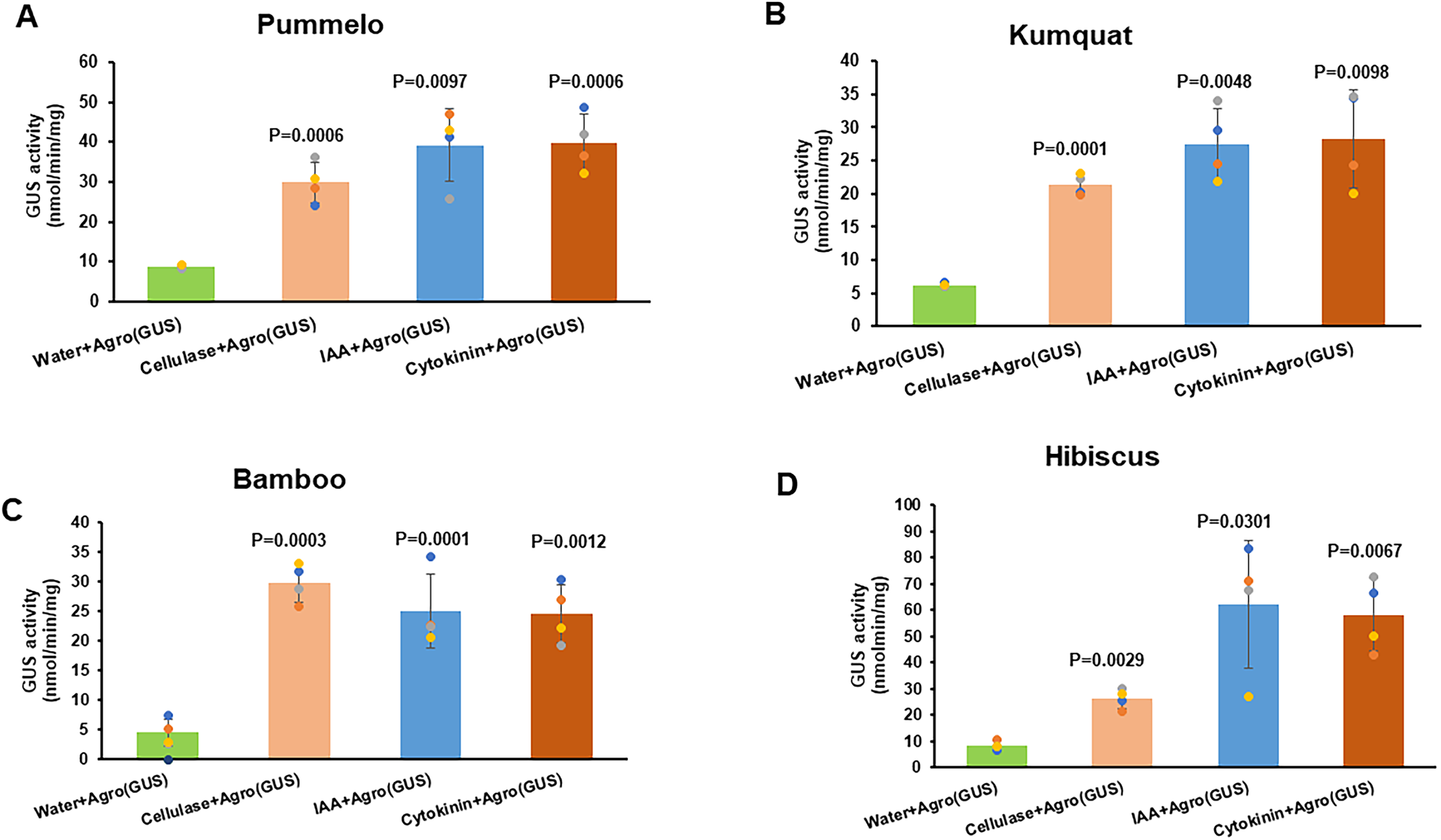
Cellulase, IAA, and cytokinin treatments enhance *Agrobacterium* mediated transient expression assay in multiple plants. A. pummelo (*C. maxima*). B. Meiwa kumquat (*Fortunella crassifolia*). C. lucky bamboo (*Dracaena sanderiana*). D. rose mallow (*Hibiscus rosa-sinensis*). *Agrobacterium tumefaciens* strain EHA105 (OD_600_=0.8) carrying the GUS binary vector was used for agroinfiltration. Leaves were first treated with cellulase enzyme (20mg/ml), IAA (20 mM), kinetin (20 mM) and followed by agroinfiltration on the same treatment site after 12 hours. Water treatment is used as control. GUS activity was measured 4 days post agroinfiltration. Error bars represent mean ± SD (n =4). All experiments were repeated twice with similar results. Student t test was conducted for statistical analysis (p<0.05).

## Discussion

In this study, we have shown that *Xcc* promotes *Agrobacterium*-mediated TE via promoting *Agrobacterium* growth, suppressing plant immune responses, triggering cell division, and upregulating plant cell wall degrading enzymes. Specifically, the first two actions enhance the competence between *Agrobacterium* and citrus. It was suggested that activation of plant immune responses is one of the main reasons for recalcitrance of *Agrobacterium*-mediated transformation of certain plant species (Rosas-Díaz *et al*., 2017). *Xcc* suppresses the expression of *PR1*, *PR2*, and *PR5* at 4 dpi, but induced *PR1* expression at 1 dpi. This expression pattern of *PR* genes is typical for compatible plant-pathogen interactions (Jones and Dangl, 2006). Our data suggest that *Xcc* suppresses salicylic acid (SA) signaling pathway at 4 dpi because *PR1*, *PR2*, and *PR5* are marker genes of SA related immune response (Hui *et al*., 1994). Our data are consistent with that expression of *NahG* which encodes SA hydroxylase to decrease SA content in planta and increases the efficiency of *Agrobacterium*-mediated TE in *Arabidopsis* (Rosas-Díaz *et al*., 2017). However, we expect that the effect of *Xcc* on plant immunity plays a minor role in promoting *Agrobacterium*-mediated TE because of the limited effect on *Agrobacterium* growth and neglectable effect on ROS levels.

*Xcc* triggered cell division especially transition through the S-phase is critical for *Xcc* promoting *Agrobacterium*-mediated TE. It was reported that active dividing cambium cells are suitable for *Agrobacterium* transformation (Peña *et al*., 2004). Citrus leaves used for transformation are fully expanded and have largely stopped cell division (Gonzalez *et al*., 2012), which is reactivated by *Xcc*. In addition, approaches that are known to promote cell division also promote *Agrobacterium*-mediated TE. For instance, overexpression of a cell cycle regulator *RepA* in maize stimulates cell division and improves maize transformation (Gordon-Kamm *et al*., 2002). Overexpression of cell cycle reprogramming proteins also increases *Agrobacterium*-mediated TE in *Nicotiana benthamiana* (Norkunas *et al*., 2018). We have demonstrated the causative relationship between cell division and *Agrobacterium*-mediated TE using cell division inhibitor mimosine. Furthermore, PthA4 is known to be responsible for causing excessive cell division of citrus leaves (Duan *et al*., 1999). Mutation of *pthA4* abolishes the promoting effect of *Xcc* on *Agrobacterium*-mediated TE, supporting the role of cell division in *Agrobacterium* transformation. *Xcc* also upregulates auxin and gibberellin response genes (Pereira *et al*., 2014) and cytokinin metabolism (Hu *et al*., 2014; Zou *et al*., 2021). Those hormones are known to promote cell division (Shimotohno *et al*., 2021). However, only auxin and cytokinin but not gibberellin promoted *Agrobacterium*-mediated TE. This might result from their different mechanisms in promoting cell division. Auxin exerts its control of cell division via transcription factors such as E2FB (Magyar *et al*., 2005); cytokinin triggers nuclear localization of a transcription factor to induce cell division (Yang *et al*., 2021); whereas gibberellin first promotes cell elongation which in turn stimulates cell division as a result of cell-growth (Sauter and Kende, 1992). Importantly, the differential effects of mimosine and colchicine on *Xcc* promoting *Agrobacterium*-mediated TE accords with that S-phase is critical in T-DNA transfer (Villemont *et al*., 1997). This is consistent with a previous report that phytohormone auxin application significantly increases the number of actively dividing citrus cells (S-phase) and results in higher transformation efficiency (Peña *et al*., 2004).

*Xcc* also promotes *Agrobacterium*-mediated TE through upregulation of cell wall degrading enzymes. We have provided direct evidence that treatment of citrus leaves with cell wall degrading enzymes improves *Agrobacterium*-mediated TE. Plant cell wall contains many phenolic compounds and sugars, which are known to trigger the *vir* genes responsible for T-DNA transfer (Cangelosi *et al*., 1990; Scott *et al*., 1985). Intriguingly, the plant cell wall degrading enzymes encoded by *Xcc* has neglectable effect on its promotion of *Agrobacterium* transformation as demonstrated by the *xpsD* mutant. *Xcc* activation of plant cell wall degrading enzymes is through PthA-LOB1 (de Souza-Neto *et al*., 2023; Su *et al*., 2023; Zhang *et al*., 2017; Zou *et al*., 2021) which likely contributes to the loss of the promoting effect of the *pthA4* mutant on *Agrobacterium*-mediated TE.

Our study has led to the discovery of practical tools to promote *Agrobacterium*-mediated TE in citrus and other plants via pretreatment with auxin, cytokinin and cellulase. Because *Xcc* is a bacterial pathogen, its application is limited in certain circumstances to avoid complicated effects of *Xcc*. *Xcc* can only be used on its citrus hosts but not on other plant species. Pretreatment with auxin, cytokinin and cellulase avoids the host range limitation of *Xcc*, thus can be used to promote *Agrobacterium*-mediated TE on other plants that are incompetent with *Agrobacterium.* Moreover, even though we only tested *Agrobacterium*-mediated TE, it is possible that cytokinin and cellulase treatment can also promote the *Agrobacterium*-mediated stable transformation as reported previously for auxin (Peña *et al*., 2004).

In sum, we demonstrate that *Xcc* promotes *Agrobacterium*-mediated TE via PthA which triggers cell division and upregulates cell wall degrading enzymes and also by improving the competence between *Agrobacterium* and citrus. Our study also identified two practical measures to improve *Agrobacterium*-mediated TE in citrus and other plants via pretreatment with plant hormones (i.e., auxin and cytokinin) and cellulase.

## Materials and Methods

### Bacterial Strains, and Bacterial Growth Conditions

The bacterial strains and plasmids used in this study are listed in supplementary Table 1. Luria-Bertani (LB) broth medium and plates were used for cultivation of *Escherichia coli* and *Agrobacterium.* Nutrient agar medium (NA) and nutrient agar (NA) plates were used for culturing *Xcc* (Sambrook *et al*. 1989). *E. coli* was grown at 37^°^C, whereas *A. tumefaciens* and *Xcc* were grown at 28^°^C. When required, growth media were supplemented with ampicillin (100 µg/ml), gentamicin (20 µg/ml), kanamycin (50 µg/ml), and rifamycin (50µg/ml).

### Pre-treatment and Agroinfiltration of Leaves

Fully expanded young citrus leaves were treated with *Xcc*, plant growth hormones (IAA, cytokinin, gibberellin) or water (control) using needleless syringes. Actively growing WT and mutant *Xcc* strains were pelleted and resuspended in sterile double distilled water (ddH2O) (5 x 10^8^ CFU/ml) before inoculation into citrus leaves using a needleless syringe. At 12 hours post treatment, agroinfiltration was performed at the same inoculation site using a needleless syringe. *A. tumefaciens* cells for agroinfiltration assay were prepared as described elsewhere (Jia and Wang, 2014). Briefly, the actively growing culture of *Agrobacterium* was pelleted and resuspended in sterile ddH_2_O to OD_600_ = 0.8. Non-treated citrus leaves were directly subjected to agroinfiltration as a non-treated control. In addition, cellulase (20 mg/ml), IAA (20 mM), and kinetin (20 mM) treatments were tested for their effect on *Agrobacterium* mediated transient expression assay in leaves of pummelo (*C. maxima*), Meiwa kumquat (*Fortunella crassifolia*), lucky bamboo (*Dracaena sanderiana*) and rose mallow (*Hibiscus rosa-sinensis*).

### Plasmid Construction

To construct the GUS vector, the fragment containing the GUS reporter gene (*uidA*) was digested with BamHI and SacI restriction enzymes from the pBI121 vector (Supplementary Table 1). The digested GUS fragment of 1812 bp was cloned into pCAMBIA1380-35S-EYFP vector (Supplementary Table 1) by replacing EYFP with GUS to make the pCAMBIA1380-35S-GUS vector (pGUS). The pGUS vector was introduced into *A. tumefaciens* strain EHA 105 by electroporation and pGUS positive transformants were selected by plating on LB agar medium supplemented with rifampicin and kanamycin. For GFP expression, the binary vector RCsVMV-erGFP-pCAMBIA-1380N-35S-BXKES-3xHA was used (Ma *et al*., 2022).

### GUS Activity Assay

The quantitative GUS assay was carried out as described previously (Hu *et al*., 2014) with modifications. Briefly, leaf discs (30 mg) were collected 4 dpi and then flash frozen in liquid nitrogen and homogenized with the Tissue Lyser II (QIAGEN, Hilden, Germany) to make fine powder at 300 rpm for 1 min. The tissue powder was mixed with 1 ml of GUS extraction buffer, vortexed vigorously, then centrifuged for 10 minutes at 4^0^C (speed ∼12000 rpm). Supernatant was collected and fluorogenic assay was performed in a plate reader (Synergy HT Plate Reader (BioTek, Winooski, VT, USA). GUS assay buffer was pre-warmed at 37°C and 25 µl of plant extract (supernatant from sample) was mixed with 225 µl of GUS assay buffer. The reaction was incubated at 37°C for 90 minutes and the reaction was stopped by adding stop solution (0.2M Na_2_CO_3_) and fluorescence was measured at 365 nm excitation and 455 nm emission wavelength. Simultaneously the 4-MU standard curve was prepared with known concentrations of 4-MU to calculate concentration of samples. Total protein concentration of samples was estimated by Bradford reagent (Bio-Rad) following the manufacturer’s protocol. GUS activity was quantified and expressed as 4MU-nmol/min/mg protein sample.

### Measurement of *Agrobacterium* Growth

*Agrobacterium* growth in *Xcc* treated and non-treated samples was measured by plate counting method. For each biological replicate, two leaf discs of 0.5 cm diameter were homogenized in sterile ddH_2_O and then plated on LB plates supplemented with rifampicin (50 µg/mL) and kanamycin (50 µg/mL) after serial dilutions. After incubation for 2 days at 28°C, the resulting colonies were counted, and *Agrobacterium* growth was analyzed.

### RNA Extraction and Quantitative RT-PCR (RT-qPCR)

Total RNA was isolated from citrus leaf samples using RNeasy plant mini kit (Qiagen, Germantown, MD, USA) following the manufacturer’s protocol. For each sample, 6-8 leaf discs from inoculated areas of three leaves from the same plant were pooled and frozen in liquid nitrogen. The samples were ground into fine powder using the Tissue Lyser II (QIAGEN, Hilden, Germany). Residual genomic DNA was removed by a TURBO DNA-free kit (Invitrogen, Carlsbad, CA, USA). cDNA was synthesized using a high-capacity cDNA reverse transcription kit (Applied Biosystems Inc., Foster City, CA). RT-qPCR was performed with Fast SYBR Green master mix (Applied Biosystems Inc., Foster City, CA) in QuantStudio 3 Real-Time PCR System (ThermoFisher, Waltham, MA). Citrus GAPDH gene was used as an endogenous control for normalization and relative gene expression was calculated by relative expression 2−^ΔΔCT^ method (Livak and Schmittgen, 2001).

### Quantification of H_2_O_2_ Concentrations

H_2_O_2_ concentrations were measured for agroinfiltrated citrus leaves with or without *Xcc* treatment (Kumar *et al*., 2011). Fresh leaf disc samples (∼0.5 g) were collected and ground in 0.1% (W/V) of trichloroacetic acid (TCA) and centrifuged at 12,000g for 15 minutes at 4°C. The collected supernatant (0.3 ml) was mixed with 1.7 ml of 1M potassium phosphate buffer (PH 7.0) and 1.0 ml of 1 M of potassium iodide (KI) solution then incubated for 5 minutes. The oxidation product was measured at Abs 390 nm a Synergy LX plate reader (BioTek, Winooski, VT, USA). H_2_O_2_ standard curve was prepared with known concentrations of H_2_O_2_ to calculate that of tested leaf samples.

## Acknowledgments

We thank Wang lab members for constructive suggestions and insightful discussions. We thank Dr. Jinyun Li for help with statistical analyses and Dr. Frank White for critical reading. This project was supported by funding from Florida Citrus Initiative Program, Citrus Research and Development Foundation 18-025, U.S. Department of Agriculture National Institute of Food and Agriculture grants 2022-70029-38471, 2021-67013-34588, 2018-70016-27412 and 2016-70016-24833, FDACS Specialty Crop Block Grant Program AM22SCBPFL1125, and Hatch project [FLA-CRC-005979] (N. Wang).

## Author Contributions

N. W. conceptualized, designed the experiments and supervised the project. T. L. performed the experiments. N. W., and T. L. wrote the manuscript.

## Figure legends

Supplementary Fig. 1. Inhibitory activity assays between *Xanthomonas citri* subsp. citri (*Xcc*) and *Agrobacterium* by cross streak method. *Agrobacterium* strain EHA 105 of OD_600_=0.8 carrying binary vector (GUS) and *Xcc* wild type strain (5X10^8^ CFU/ml) were used for this test. Nutrient agar plates without any antibiotics were used for the test. 20 μL of *Xcc* strain was streaked vertically and 10 μL of *Agrobacterium* was streaked perpendicular to the Xcc strain then incubated for 2 days at 28^0^C and photos were taken. Experiment was repeated twice with similar results. No inhibitory effect was observed.

Supplementary Fig. 2. ROS levels in citrus leaves in agroinfiltration assay. H_2_O_2_ concentrations were quantified at different time points indicated above. Citrus leaves were treated with Xcc of 5X10^8^ CFU/ml and sterile water, followed by agroinfiltration after 12 hours. Direct agroinfiltration was performed for control purpose. Values represent means ± SD (n=4). Student’s *t* test was used for statistical analysis. Same letter indicate there is no significant differences (P-value > 0.05). All experiments were repeated twice with similar results.

Supplementary Fig. 3. Transient GFP expression in citrus leaves facilitated by *Xcc*. Citrus leaves were treated with Xcc WT (5X10^8^ CFU/ml) followed by agroinfiltration on the same treatment site 12 hours later. GFP observation was conducted at 4 dpi. Agro: Agroinfiltration; DPI: days post agroinfiltration; *Xcc*: *Xanthomonas citri* subsp. *citri*; WT: wild type; ΔgumC: *gumC* mutant of Xcc; ΔxpsD: *xpsD* mutant of *Xcc*; XccpthA4:Tn5: *pthA4* Tn5 insertion mutant of *Xcc*. Water treatment, agroinfiltration alone, and leaf alone were used as negative controls. GFP expression was detected by confocal microscopy.

Supplementary Fig. 4. PthA4 is required for the *Xcc-facilitated* agroinfiltration. A. GUS assay. Citrus leaves were treated with *Xcc* strains followed by agroinfiltration on same treatment site after 12 hours. Water treatment and agroinfiltration alone were used as negative controls. *Agrobacterium tumefaciens* strain EHA 105 carrying GUS binary vector was used for agroinfiltration. GUS assay was performed at 4 dpi. The values represent mean ± SD (n=4). Different letters indicate significant difference (P<0.05, One-way ANOVA with post-hoc Tukey Test). All experiments were repeated twice with similar results. B. Histochemical GUS staining of citrus leaf tissues in A. Agro: Agroinfiltration; DPI: days post agroinfiltration; *Xcc*: *Xanthomonas citri* subsp. *citri*; WT: wild type; *pthA4* mutant: *pthA4:*Tn5 insertion mutant of *Xcc*; *XccpthA4* complementation: *pthA4:*Tn5 insertion mutant of *Xcc* complemented with wild type *pthA4*.

Supplementary Fig. 5. Gene expression of a citrus endoglucanase gene in agroinfiltration assay of citrus. Citrus leaves were treated with 5X10^8^ CFU/ml of Xcc strains (WT, *pthA4* mutant and *pthA4* mutant complementation strain) followed by agroinfiltration 12 hours after treatment. Agroinfiltration was performed with *Agrobacterium tumefaciens* strain EHA 105 carrying the GUS binary vector. Agroinfiltration without any treatment was performed as a negative control (Agro). Pretreatment with water was used as the mock inoculation control. RT-qPCR was conducted 4 days past agroinfiltration. Citrus *GAPDH* gene was used as internal control for normalization. Error bars indicate mean ±SD (n=4). One way ANOVA followed by Tukey HSD test was conducted for statistical analysis. Different letters indicate significant statistical difference (p<0.05).

**Supplementary Table 1. Bacterial strains and plasmids used in this study**

